# Regeneration-associated utrophin expression in skeletal muscle: implications for utrophin-targeted therapies

**DOI:** 10.64898/2026.09.10.750746

**Authors:** Simon Guiraud, Benjamin Edwards, Sarah E. Squire, Kay E. Davies

**Affiliations:** MDUK Neuromuscular Centre at the University of Oxford, Department of Physiology, Anatomy and Genetics, Oxford OX1 3PT, United Kingdom

**Keywords:** Duchenne muscular dystrophy, utrophin, regeneration, biomarkers, embryonic myosin

## Abstract

Duchenne muscular dystrophy (DMD) is a lethal, X-linked muscle-wasting disease caused by loss of dystrophin. Utrophin, a structural and functional paralogue of dystrophin, can compensate for dystrophin deficiency and represents a therapeutic target applicable to patients irrespective of their DMD mutation. However, utrophin is also naturally upregulated at the sarcolemma of regenerating myofibres, creating an important confounding factor when utrophin levels are used to evaluate therapeutic activity. Here, we defined the temporal relationship between utrophin expression and complementary markers of skeletal muscle regeneration in wild-type and dystrophic mdx muscle following cardiotoxin-induced injury. Utrophin protein and mRNA dynamics were assessed together with dystrophin, embryonic myosin (MyHC-emb), centronucleation, cross-sectional area (CSA), and minimal Feret’s diameter (MinFeret). MyHC-emb and utrophin were strongly associated with the early regenerative phase, although MyHC-emb peaked before maximal utrophin protein expression. In contrast, CSA and MinFeret showed an inverse temporal relationship with utrophin, while centronucleation persisted after active regeneration and did not discriminate early from late regenerative stages. Utrophin also remained elevated at the dystrophic sarcolemma for longer than in wild-type muscle. These findings demonstrate that utrophin quantification cannot be interpreted independently of the regenerative state of muscle. Combining MyHC-emb with morphological indices such as CSA and MinFeret provides a complementary framework for contextualizing regeneration-associated utrophin and should improve interpretation of utrophin changes in preclinical evaluation of utrophin-targeted therapies.

## Introduction

Duchenne Muscular Dystrophy (DMD) is a fatal X-linked neuromuscular disorder affecting 1 in 5000 newborn males [1, 2] making it one of the most common recessive diseases in the human population. Affected boys are generally diagnosed between 2 and 5 years of age and develop progressive muscle degeneration leading to orthopaedic, respiratory and cardiac complications. Improvements in multidisciplinary care have extended survival, although life expectancy remains substantially reduced [3, 51]. At the molecular level, DMD is caused by loss of function mutations in the dystrophin gene (DMD, MIM #310200) [4, 5]. The dystrophin gene is the largest in the human genome (2.3 megabases) and presents the highest de novo mutation rate, predominantly resulting in deletions of the gene [6]. The 14 kb RNA transcript is expressed principally in muscle and translated into a 427 kDa cytoskeletal dystrophin protein, critical to maintain fibre strength, flexibility and stability in skeletal muscle. Dystrophin establishes a mechanical link between the extracellular matrix and the intracellular cytoskeletal actin in myofibres through the dystrophin-associated protein complex (DAPC) [7] and acts as a molecular shock absorber during muscle contraction. At the cellular level, dystrophin deficiency in DMD or reduction in the milder Becker muscular dystrophy (BMD, MIM #300376; [8]) leads to costamere perturbation, sarcolemma fragility and subsequent chronic inflammation associated with rounds of degeneration and regeneration. This subsequently leads to muscle necrosis, fibrotic events and ultimately loss of muscle fibres with reduction in muscle mass and function.

There is currently no cure for DMD. Available therapies can slow disease progression or address specific disease mechanisms, but clinical benefit remains incomplete, and parallel efforts continue to develop gene-based, cell-based and pharmacological strategies [9, 10, 52]. A promising approach, applicable to all DMD patients irrespective of their dystrophin mutation, is to upregulate utrophin, a structural and functional autosomal paralogue of dystrophin [11, 53]. Despite subtle differences [12–14], utrophin and dystrophin share a high level of structural identity and a similar intron-exon structure [15, 16]. Pre-clinical studies in murine and dog models of DMD demonstrated that truncated utrophin mini and micro-genes [17–19] partially prevent the pathology. Transgenic expression of the full length utrophin gene in the mdx mouse model of DMD [20] suppresses functional signs of dystrophinopathy in a dose dependent way [21] without toxicity [22]. Whereas dystrophin is mainly confined to adult skeletal, smooth, cardiac muscles and brain [23], utrophin is ubiquitously localised at the sarcolemma in early [24] developing muscles [25] and progressively replaced by dystrophin towards birth [26] to be limited to the neuromuscular (NMJ), the myotendinous (MTJ) junctions in adult muscles [27] and blood vessels [28]. Importantly, in mdx muscle, utrophin is increased as part of the repair process by ∼2-3-fold in muscle and localised at the sarcolemma of regenerating myofibres [29]. A respective ∼2.5 and ∼4-fold utrophin up-regulation was also described in BMD and DMD patients [30]. A small increase in utrophin was previously reported to delay the age when DMD patients are wheelchair bound [31] although other studies described no relationship [32] or a counterintuitive negative correlation of utrophin expression with clinical severity [33]. The level of regeneration and the subsequent level of utrophin vary between muscle type [34] and are age dependent in both animals and DMD patients [26, 35]. Therefore, utrophin, often described as a regeneration-associated protein [36], can only be accurately described when correlated with regeneration markers [37]. Dystrophic skeletal muscles are characterised by excessive fibre size variation, large rounded hypertrophic fibres, fibres with central nuclei, as well as hypercontracted fibres and clusters of small regenerating myofibres. These histological characteristics offer several indices of skeletal muscle regeneration, a hallmark of the disease. The most commonly used markers of regeneration are the presence of centronucleated muscle fibres (CNF) (Treat-NMD SOP DMD_M.1.2.001) and a morphological change in size of nascent muscle cells determined by cross-sectional area (CSA) and minimal Feret’s diameter (MinFeret) (Treat-NMD SOP DMD_M.1.2.001). The presence of embryonic myosin (MyHC-emb) in dystrophic muscles [38] is a meaningful indicator of muscle damage which correlates with functional motor score in BMD and DMD patients [39]. Our recent study demonstrated that MyHC-emb is a robust marker of regeneration at different ages and in different muscles of the mdx mice [35]. Nevertheless, there is no single marker that unequivocally identifies a regenerating fibre and the use of a panel of complementary biomarkers will provide a more accurate and complete view on the regenerative status of the muscle. Recent transcriptomic and histological studies have also identified additional regeneration-associated markers in human dystrophinopathies, further supporting a multimarker approach [54].

In the present study, we investigated the temporal relationship between utrophin expression and skeletal muscle regeneration in wt and mdx muscle using a time course of cardiotoxin (CTX)-induced injury. Because endogenous utrophin is itself induced during muscle repair, changes in total or sarcolemmal utrophin may be difficult to attribute to a therapeutic intervention without simultaneously defining the regenerative state of the tissue. We therefore assessed utrophin and dystrophin localization and expression together with complementary regenerative markers, including centronucleation, CSA, MinFeret, and MyHC-emb. Our aim was not simply to identify markers associated with regeneration, but to determine which combination of measurements best contextualizes regeneration-associated utrophin across the regenerative time course. This framework is intended to improve interpretation of utrophin measurements in preclinical studies of utrophin-targeted therapies.

## Results

### Time course of CTX-induced muscle regeneration and associated regenerative markers

In 7-week-old wild-type (wt) skeletal muscle, fibres display a uniform morphology with intact sarcoplasm and peripheral nuclei (Fig. 1A). By contrast, *mdx* muscles at the same age show pronounced fibre size variability and central nuclei, reflecting the ongoing cycles of degeneration and regeneration that follow the DMD-like crisis occurring between 3 and 5 weeks of age [41].

**Fig. 1.**
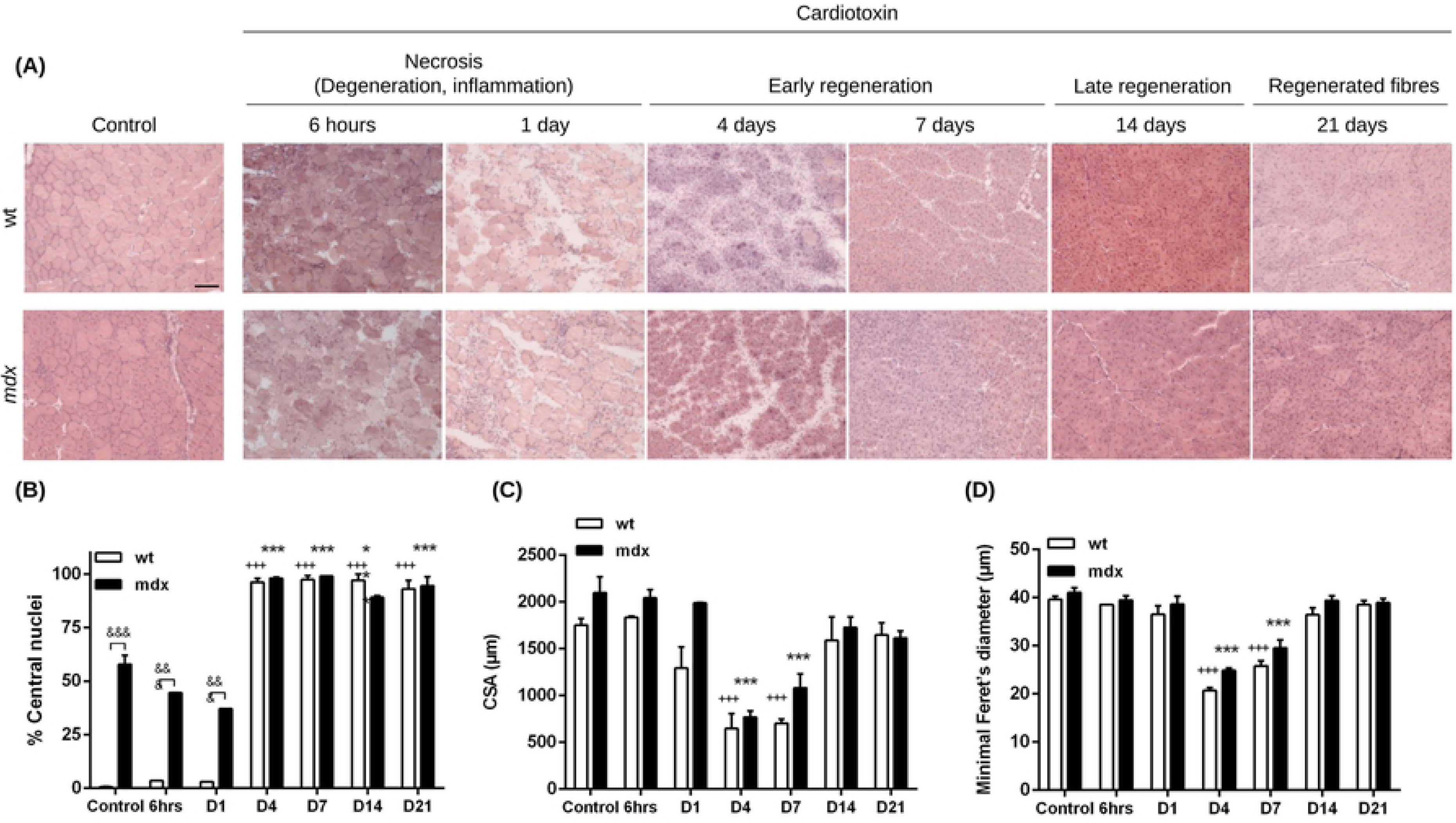
Morphological parameters following cardiotoxin injection in wt and mdx skeletal muscle. (A) Representative H&E-stained transverse sections of tibialis anterior (TA) muscles from wt and mdx mice after a single intramuscular injection, collected at different time points (6 h, 1, 4, 7, 14, and 21 days). Scale bar: 100 µm. (B) Percentage of centrally nucleated fibers (CNFs) in saline-injected controls and CTX-injected wt and mdx groups. (C) Cross-sectional area (CSA) in control and CTX-injected wt and mdx groups. (D) Minimal Feret’s diameter in control and CTX-injected wt and mdx groups. Values are expressed as mean ± SEM (n = 5 per condition). *p<0.05, **p<0.01, ***p<0.001 mdx relative to control mdx, +p<0.05, ++p<0.01, +++p<0.001 wt relative to control wt, &p<0.05, &&p<0.01, &&&p<0.001 mdx compared to wt.

To investigate muscle regeneration in both wt and *mdx* muscle, we performed a single intramuscular injection of CTX into the tibialis anterior (TA) of 4-7-week-old animals and examined histological and morphological changes over time (Fig. 1A). At 6 h and 1-day post-CTX, extensive fibre necrosis and inflammatory infiltration were observed in both genotypes. Necrotic fibres exhibited irregular contours and fragmented sarcoplasm. This initial inflammatory and acute degenerative necrotic phase is rapidly followed at day 4-7 by an early regeneration characterised by numerous small fibres with central nuclei. By 14–21 days post-injection, fibres had enlarged to near control size, and muscle architecture appeared largely restored except for persistent central nuclei.

We next studied several indices of regeneration. CNF is a commonly used marker of regeneration [21, 42]. As expected, prior to the CTX injection, CNF is negligible in wt muscle whereas ∼55 % of mdx fibres were centrally nucleated (Fig. 1B), reflecting ongoing regeneration in dystrophic muscles. During the necrotic phase (6 h–1-day post-CTX), CNF proportion transiently decreased due to fibre loss. From day 4 post-CTX injection, the majority of myofibres are centrally nucleated. The high level of CNF is maintained throughout day 7, 14 and 21. Consequently, CNF alone does not discriminate between early and late regeneration stages. In contrast, the CSA and MinFeret provided clearer indicators of regenerative progression. CSA significantly decreased during the early regeneration phase (days 4–7; Fig. 1C) and returned to baseline levels by days 14–21. The MinFeret parameter, taking into account the orientation of the sectioning angle, showed a similar temporal pattern (Fig. 1D).

Together, these data show that CSA and MinFeret track the progression from early regeneration toward restoration of fibre size, whereas CNF remains elevated after the active regenerative phase. Thus, centronucleation is useful as a cumulative indicator of previous degeneration/regeneration but, when used alone, provides limited temporal information for interpreting regeneration-associated changes in utrophin.

### Dystrophin, utrophin, and MyHC-emb expression in CTX-induced regenerating wt muscle

In healthy muscle, dystrophin localises to the sarcolemma (Fig. 2A), whereas utrophin is restricted to neuromuscular and myotendinous junctions and to blood vessels (Fig. 3A). MyHC-emb is undetectable in control muscle by immunofluorescence (Figs. 2A and 3A) and quantitative analysis (Fig. 4), and is also undetectable by Western blot (Fig. 5). During the necrotic phase, dystrophin and utrophin protein levels markedly decreased relative to untreated wt muscle (6 h: DYS, 0.10-fold; UTR, 0.58-fold; day 1: DYS, 0.10-fold; UTR, 0.58-fold; Figs. 2, 3 and 5) while MyHC-emb remained negligible. At 4 days post-CTX, dystrophin expression remained low (4-day: 0.02-fold), whereas utrophin was strongly upregulated at the sarcolemma 4 and 7 days post-CTX injection (4 day: 2.6-fold; UTR 7 day: 4.7-fold). This increase coincided with abundant MyHC-emb expression (83.8 % positive fibres; 444× increase in protein vs. control; Figs. 2–5). Utrophin and MyHC-emb thus peaked at days 7 and 4 respectively (Fig. 5A, D and F). CTX administration profoundly alters the total protein profile and none of the usual normalisation proteins such as vinculin, a-actinin and GAPDH provide a stable and consistent protein level across the time course of CTX-induced muscle regeneration (S1 Fig). Therefore, a non-relative quantification of dystrophin, utrophin and MyHC-emb protein was used to avoid bias. High levels of utrophin and MyHC-emb mark the early stage of the regenerative processes (days 4-7), sarcolemmal utrophin expression gradually returned to baseline (days 14–21) and MyHC-emb became undetectable. Dystrophin reappeared from day 7 onward, reaching control levels and recapitulating its developmental expression pattern (Figs. 2A, 5A–C).

**Fig. 2.**
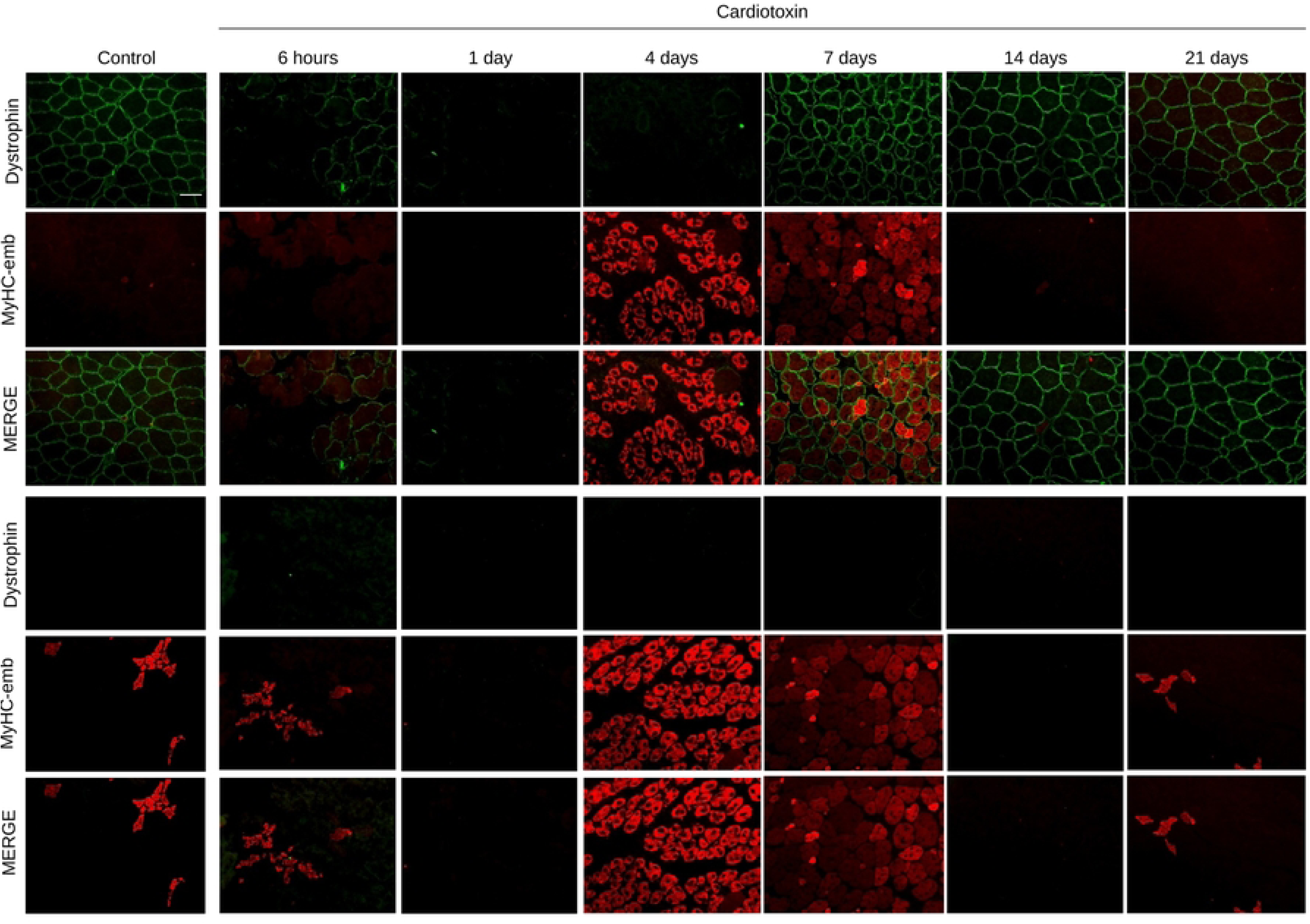
Localization of dystrophin and MyHC-emb in wt and mdx TA muscle after cardiotoxin injection. (A) Representative co-immunofluorescence images of dystrophin and MyHC-emb in control and CTX-injected wt mice across the time course of CTX-induced muscle regeneration. (B) Dystrophin and MyHC-emb localization in control and CTX-injected mdx mice. n = 5 per condition. Scale bar: 100 µm.

**Fig. 3.**
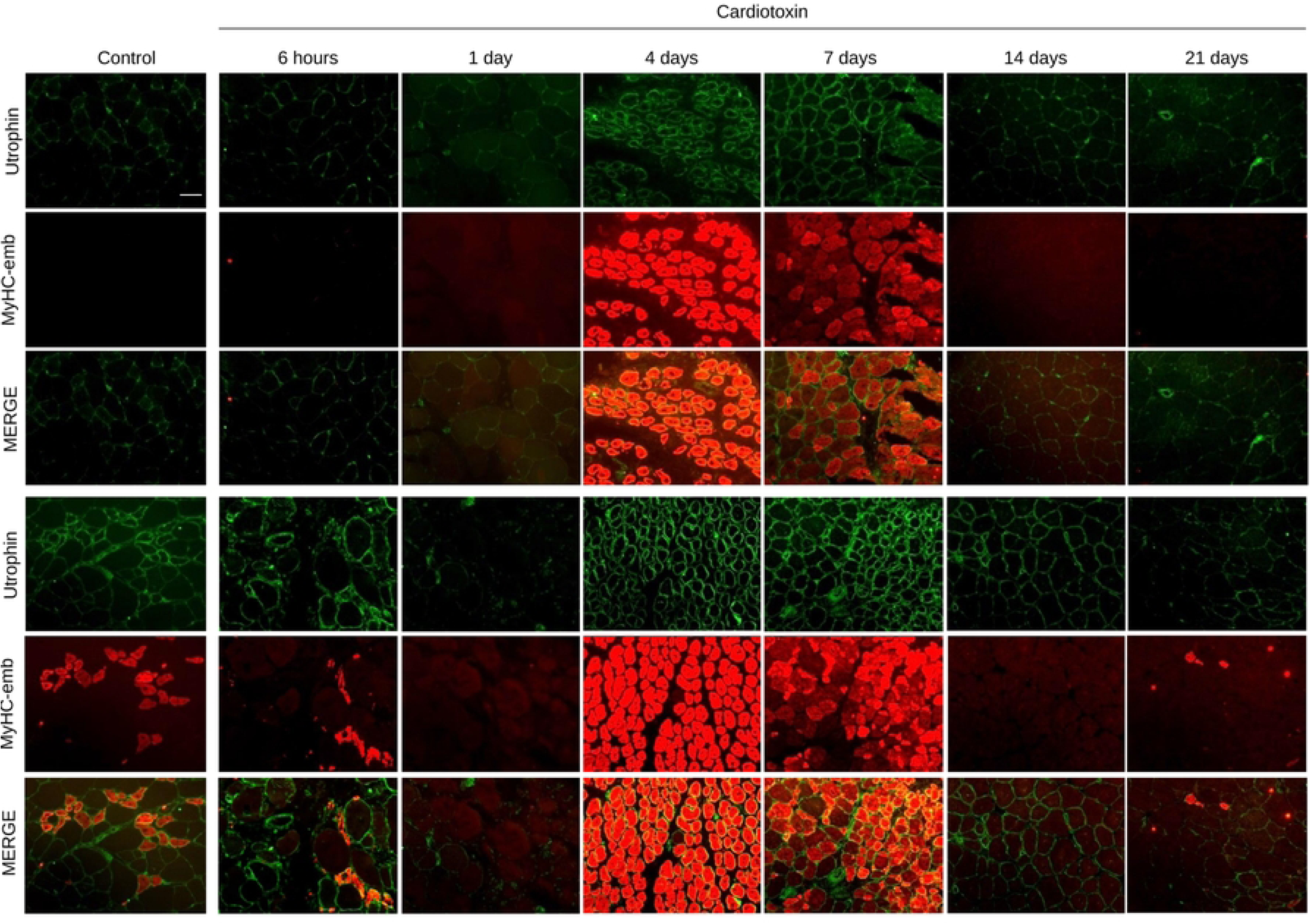
Localization of utrophin and MyHC-emb in wt and mdx TA muscle after cardiotoxin injection. (A) Representative co-immunofluorescence images of utrophin and MyHC-emb in control and CTX-injected wt mice across the time course of CTX-induced muscle regeneration. (B) Utrophin and MyHC-emb localization in control and CTX-injected mdx mice. n = 5 per condition. Scale bar: 100 µm.

**Fig. 4.**
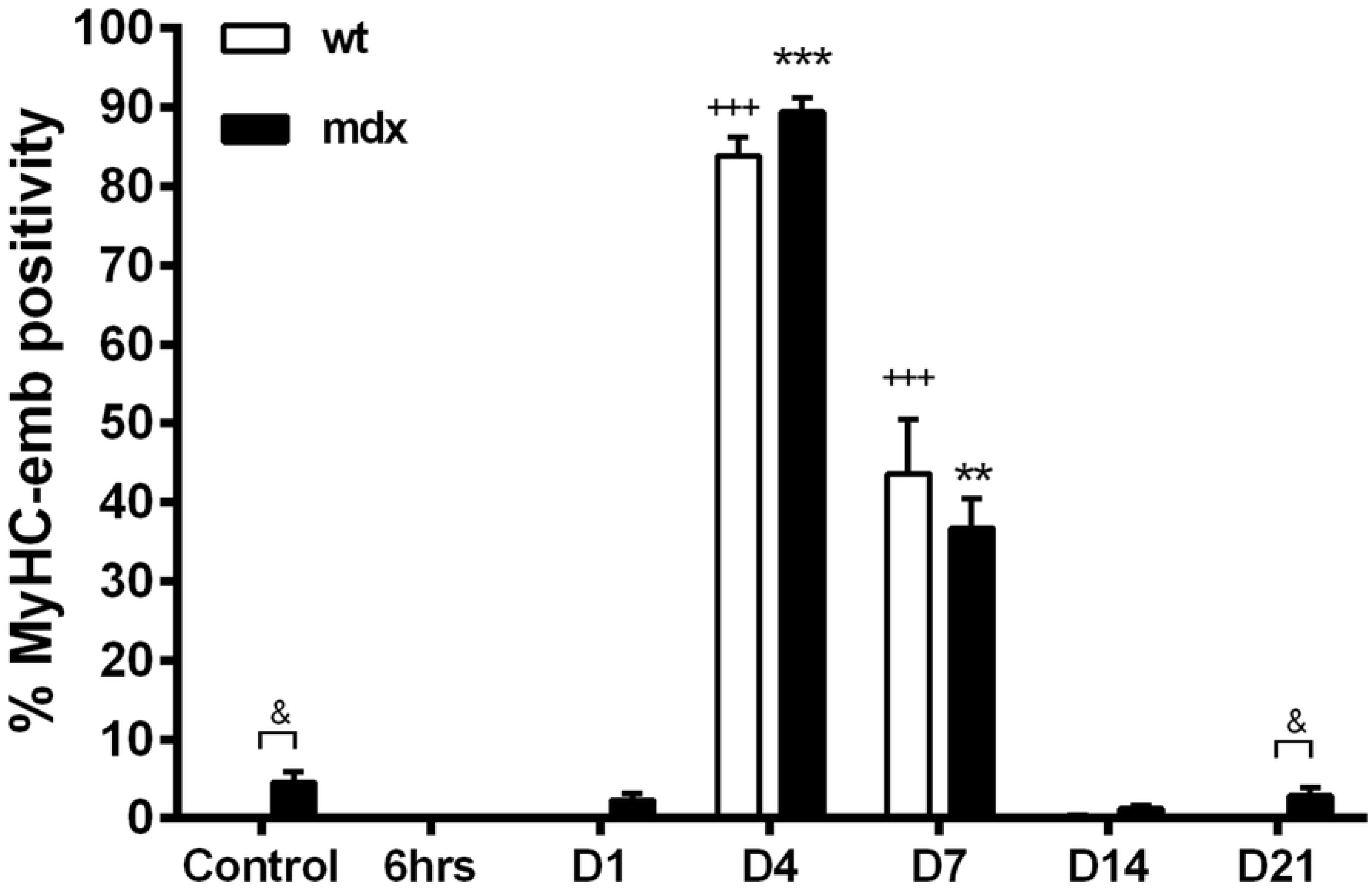
MyHC-emb levels in wt and mdx muscle. Quantification of MyHC-emb–positive myofibers in control and CTX-injected wt and mdx mice across the time course of CTX-induced muscle regeneration. Values are expressed as mean ± SEM (n = 5 per condition). *p<0.05, **p<0.01, ***p<0.001 vs mdx, +p<0.05, ++p<0.01, +++p<0.001 vs wt control, &p<0.05, &&p<0.01, &&&p<0.001 mdx vs wt.

**Fig. 5.**
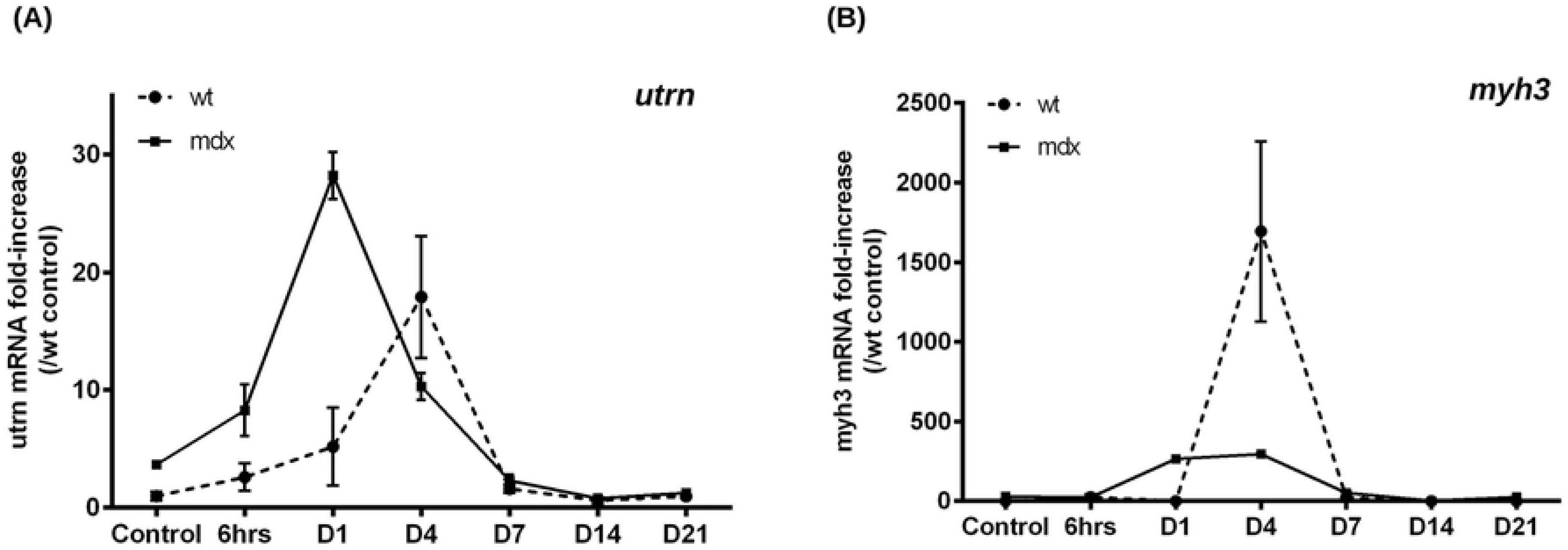
Dystrophin, utrophin, and MyHC-emb protein levels in wt and mdx muscle following CTX injection. (A) Immunoblot analysis of dystrophin, utrophin, and MyHC-emb in individual control and CTX-injected wt TA muscles. (B) Immunoblot analysis of dystrophin, utrophin, and MyHC-emb in individual control and CTX-injected mdx TA muscles. (C) Dystrophin protein levels in wt mice relative to wt controls. (D) Utrophin protein levels in wt mice relative to wt controls. (E) Utrophin protein levels in mdx mice relative to wt controls. (F) MyHC-emb protein levels in wt mice relative to wt controls. (G) MyHC-emb protein levels in mdx mice relative to wt controls. Values are expressed as mean ± SEM (n = 5 per condition). *p < 0.05, **p < 0.01, ***p < 0.001 vs mdx control; +p < 0.05, ++p < 0.01, +++p < 0.001 vs wt control; &p<0.05, &&p<0.01, &&&p<0.001 mdx vs wt.

At the transcript level, utrn and myh3 mRNA expression (normalised to the stable housekeeping gene gapdh; S2 Fig) paralleled protein dynamics. Both transcripts peaked at day 4 (17.9× and 1693× over control for utrn and myh3, respectively; Fig. 6A and B). Interestingly, utrn mRNA was transiently elevated as early as day 1 (5.2×), consistent with early transcriptional activation.

**Fig. 6.**
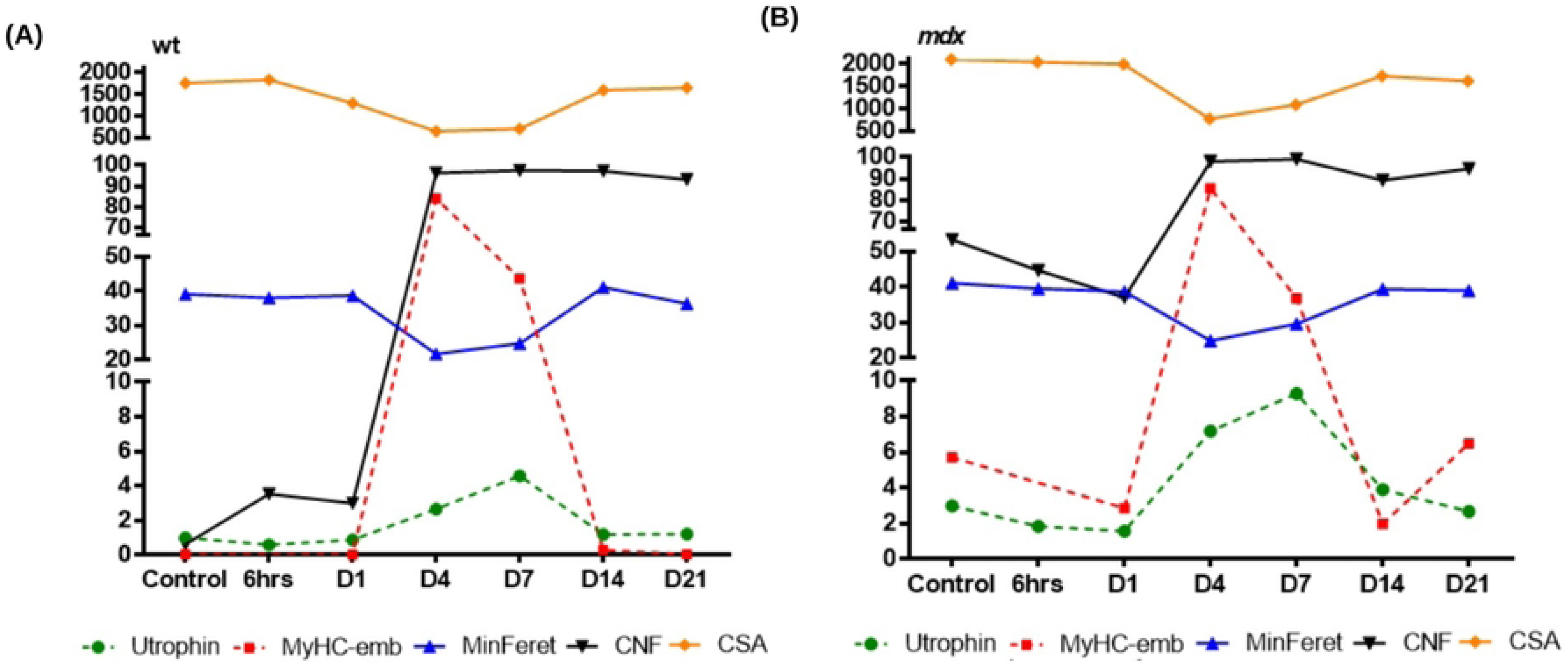
Utrophin and Myh3 mRNA expression during CTX-induced muscle regeneration. (A) utrn mRNA levels normalized to gapdh in control and CTX-injected wt and mdx TA muscles. (B) Myh3 mRNA levels normalized to gapdh in control and CTX-injected wt and mdx TA muscles.

Collectively, these data demonstrate a close temporal relationship between MyHC-emb and regeneration-associated utrophin following CTX injury, while also revealing a temporal offset between the two proteins. MyHC-emb therefore provides a sensitive marker of active regeneration that can be used alongside morphological parameters to contextualize changes in utrophin expression.

### Utrophin and MyHC-emb expression in CTX-induced regenerating *mdx* muscle

Dystrophic *mdx* muscle undergoes prolonged and repeated bouts of degeneration and regeneration between 3-10 weeks of age. At baseline, utrophin protein levels were ∼3× higher than in wt controls (Figs. 5D–E), localising to the sarcolemma and co-expressed with MyHC-emb–positive regenerating fibres (Figs. 3B, 4).

During the necrotic phase (6 h–1-day post-CTX), utrophin levels decreased to 0.61× and 0.51× relative to untreated mdx muscle (Figs. 3B, 5B, E). Subsequently, utrophin increased during regeneration, reaching 2.4× and 3.1× control levels at days 4 and 7, respectively (Figs. 5B, E). This utrophin rise paralleled high MyHC-emb expression (89.4 % positive fibres at day 4; 26 % at day 7; Figs. 4, 5B and G). From day 14 to 21, utrophin levels gradually declined but unlike wt, remained slightly elevated compared to mdx controls, with homogeneous sarcolemmal distribution (Figs. 3B, 5B, E).

Quantitative PCR confirmed significantly higher basal *utrn* (3.7×) and *myh3* (29×) transcript levels in *mdx* compared to wt muscle (Figs. 6A and B). Following CTX injection, *utrn* mRNA increased early - 8.3× (6 h), 28.2× (1 day), and 10× (4 days) - suggesting an accelerated transcriptional response (Fig. 6A). *Myh3* transcripts peaked between 1 and 7 days (265× (day 1), 295× (day 4), and 53× (day 7), Fig. 6B).

Overall, these results show that utrophin and MyHC-emb are coordinately induced during regeneration in dystrophic muscle following CTX injection, with earlier utrn transcriptional activation in mdx than in wt muscle. The persistence of utrophin after MyHC-emb declines further indicates that no single regenerative marker is sufficient to define the origin of an increased utrophin signal.

## Discussion

Utrophin is a regeneration-associated protein that reappears at the sarcolemma during muscle repair. This creates a specific challenge for the development of utrophin-targeted therapies: an increase in utrophin measured in dystrophic muscle may reflect pharmacological target engagement, endogenous regeneration, or a combination of both. Accurate interpretation of utrophin expression therefore requires simultaneous assessment of the regenerative state of the tissue. In this study, we used CTX-induced regeneration in wt and mdx skeletal muscle to define the temporal relationship between utrophin and complementary regenerative markers and to establish a practical framework for interpreting regeneration-associated utrophin.

In this study, cardiotoxin (CTX) was used to induce skeletal muscle regeneration [40], generating reproducible histological patterns of injury and repair [45] that recapitulate myogenic events in wt and mdx muscles [26]. As illustrated in Fig. 7, utrophin protein levels increased in temporal association with MyHC-emb expression. However, MyHC-emb peaked four days post-CTX injection, whereas utrophin levels reached a maximum at seven days. The temporal delay between MyHC-emb and utrophin accumulation may reflect differences in transcriptional, translational and/or protein-stability kinetics. These findings highlight a temporal shift between utrophin and MyHC-emb expression and emphasize the need for a panel of complementary regenerative markers to fully capture utrophin dynamics during regeneration.

**Fig. 7.** Temporal relationship between utrophin protein levels and key regenerative markers. (A) Temporal profiles of utrophin protein levels (fold change), MyHC-emb (%), minimal Feret’s diameter (µm), CSA (µm²), and CNF (%) across the time course of CTX-induced muscle regeneration in wt skeletal muscle. (B) Corresponding temporal profiles of utrophin and regenerative markers across the same time course in mdx skeletal muscle.

Our data indicate that MyHC-emb, CSA, and MinFeret provide complementary information for this purpose. MyHC-emb closely marks the active regenerative phase and rises before maximal utrophin protein accumulation, whereas CSA and MinFeret decrease as small regenerating fibres emerge and recover as fibre size is restored. By contrast, central nuclei persist after the active regenerative phase and therefore provide limited temporal resolution when used alone. Importantly, these measurements can be obtained from the same muscle biopsy and, in combination, can help determine whether an elevated utrophin signal occurs in a strongly regenerative environment or in fibres with limited evidence of active regeneration. This distinction is particularly relevant for utrophin-targeted therapies, which are expected both to increase sarcolemmal utrophin and, if effective, to reduce ongoing degeneration and regeneration.

The goal of utrophin-based therapies for DMD is to localise and maintain utrophin signal at the sarcolemma of adult myofibres [11]. Recent studies using distinct genetic and pharmacological approaches further support the therapeutic potential of increasing endogenous utrophin expression [55–57]. Utrophin therapies will reduce regeneration processes and consequently regeneration-associated utrophin levels in dystrophic muscle [21, 37, 42]. Hence, the use of regenerative biomarkers, applicable to all utrophin strategies in both preclinical and clinical settings, is essential to assess the extent to which changes in utrophin expression are associated with regeneration or therapeutic intervention. Our data demonstrated that CNF quantification is not suitable to monitor the utrophin signal in this context, as regenerating fibres may retain central nuclei for unclear reasons. Previous reports postulate that CNF may reflect a compensatory activation of defective myonuclei in DMD rather than regeneration per se [47]. Nonetheless, the prevalence of centronucleation remains a useful cumulative index of prior fibre necrosis and subsequent regenerative events, and may still reveal therapeutic benefit when treatment is initiated before the mdx crisis (2–3 weeks of age) [42, 43]. Additional regulators of muscle regeneration – such as long non-coding RNAs and conserved microRNAs (miR) – could complement conventional regenerative indicators, providing a more complete picture of regenerative processes [37].

The present study was designed to define these relationships in a controlled experimental setting using CTX-induced regeneration of mouse TA muscle. CTX provides a synchronized and reproducible injury-response model, but it does not reproduce the full complexity or chronicity of human DMD muscle pathology. The proposed marker framework should therefore be viewed as a basis for interpreting preclinical utrophin measurements rather than as a validated classifier of therapeutic versus endogenous utrophin. Validation in human DMD biopsies, and particularly in longitudinal samples obtained during therapeutic intervention, will be required to establish its clinical utility.

Across the CTX-induced regeneration time course, utrophin, MyHC-emb, CSA, and MinFeret profiles were globally similar in wt and *mdx* muscles, but with key differences. As expected, total utrophin protein was threefold higher in dystrophic muscle compared to wt and localized at the sarcolemma of regenerating myofibres. Following early regeneration events (4–7 days post-CTX), utrophin level returned to baseline after 14 days in wt but remained elevated in *mdx* muscle. Thus, utrophin persists at the sarcolemma for approximately 7 days in wt but for about 14 days in dystrophic muscle – possibly reflecting increased utrophin stability at the dystrophic sarcolemma in the absence of dystrophin. Post-transcriptional regulatory mechanisms are known to modulate utrophin levels in skeletal myofibres [48], and stabilization of the protein at the dystrophic sarcolemma may further contribute to this persistence. Interestingly, contrary to earlier reports of discordance between utrophin protein and transcript levels [48], we observed a significant 30-fold induction of *utrn* mRNA preceding protein accumulation, suggesting transcriptional activation of the utrophin promoter in dystrophic muscle. Following CTX treatment, utrophin increased 4.6-fold in wt and 3.1-fold in *mdx* compared with their respective untreated controls. The stronger regenerative capacity of wt muscle, reflected by higher MyHC-emb expression, may explain this difference. Interestingly, the maximal utrophin increase in *mdx* muscle was comparable to UTRN level in transgenic *mdx*-Fiona mice [35, 49, 50], suggesting that a 4–5-fold elevation above wt levels may approximate the range of maximal utrophin expression achievable in these experimental models.

Dystrophin and utrophin are expressed in an apparent reciprocal manner [26]. However, we recently demonstrated that both can be co-expressed and co-localized at the same sarcolemma, and that utrophin and dystrophin combination therapies may yield additive therapeutic benefits [50]. The present CTX-induced regeneration study confirms that utrophin and dystrophin can be respectively expressed, at high (4.6-fold increase) and normal (0.93 of the wt dystrophin) levels at the same muscle membrane. These findings reinforce the potential of utrophin-based approaches for all Becker muscular dystrophy (BMD) patients and for DMD patients treated with exon-skipping approaches and potentially with micro-dystrophin gene therapy.

In summary, our results show that utrophin expression cannot be interpreted independently of skeletal muscle regeneration. MyHC-emb provides a sensitive marker of active regeneration, while CSA and MinFeret add complementary morphological information and CNF primarily reflects the cumulative history of regenerative events. The combined assessment of utrophin with MyHC-emb, CSA, and MinFeret therefore provides a practical framework for contextualizing regeneration-associated utrophin in muscle. Applying such a framework should improve the interpretation of utrophin changes in preclinical studies of utrophin-targeted therapies and provides a rationale for future validation in DMD clinical biopsy samples.

## Materials and Methods

### Ethics Statement

All animal procedures complied with the UK Home Office regulations, in accordance with the European Community Directive 86/609/EEC (1986). Experiments were conducted under Certificate of Designation number XEC303F12 and Project Licence number 30/3104, following approval by the Joint Departmental Ethics Review Committee of the University of Oxford Departments of Physiology, Anatomy & Genetics and Experimental Psychology.

### Mice

Wild-type C57BL/10ScSnOlaHsd (C57BL/10), dystrophin-deficient C57BL/10ScSn-*Dmdmdx*/J (*mdx*). C57BL/10 mice were obtained from Envigo (UK) and all other mouse strains were bred in the Biomedical Services facility, University of Oxford.

### Cardiotoxin preparation and injection

Male C57BL/10 and mdx mice were 4–7 weeks old at the time of CTX administration, with age at injection adjusted according to the experimental time point. Mice were anesthetized with isoflurane, and 100 µL of cardiotoxin (10 µM in saline; LATOXAN) was injected into the right tibialis anterior (TA) muscle using a 30-gauge needle. The contralateral TA muscle received 0.9% saline as a control. Muscles were harvested at defined time points post-injection to ensure all mice were 7 weeks old at the time of collection.

### Histology

Frozen TA sections (10 µm) were air-dried for 10 min, stained in hematoxylin solution (Sigma, GHS232) for 8 min, rinsed in tap water, and differentiated in 70% ethanol containing 1% HCl for 10 s. Sections were then stained in 1% Eosin Y (Sigma, E4382) in 80% ethanol for 5 s, dehydrated through graded alcohols, cleared in Histochoice (Sigma, H2779), and mounted with Histomount (National Diagnostics, HS-103). Images were acquired using an Axioplan 2 Microscope System (Carl Zeiss, Germany).

### Immunofluorescence Histology

Frozen TA sections (10 µm) were fixed in acetone for 10 min, washed in PBS, and blocked for 1 h using the M.O.M.™ Blocking Kit (FMK-2202, Vector Laboratories). After two 2-min washes, sections were incubated overnight at 4 °C with primary antibodies: goat polyclonal anti-utrophin (1:500, URD40), rabbit monoclonal anti-dystrophin (1:2000, ab15277, Abcam), rat monoclonal anti-laminin-α2 (1:100, sc-59854, Santa Cruz Biotechnology), and directly conjugated anti-MYH3 (F1.652) Alexa Fluor® 594 (1:100, sc-53091 AF594, Santa Cruz Biotechnology). Following additional washes and a 5 min incubation in M.O.M. diluent, sections were incubated for 2 h at room temperature with anti-goat Alexa Fluor® 488 (1:2000, A11055, Life Technologies) and anti-rat Alexa Fluor® 488 (1:2000, A11006, Life Technologies), as appropriate. Fluorescent images were obtained using an Axioplan 2 Microscope System (Carl Zeiss, Germany) equipped with a multi-acquisition module.

### Embryonic myosin quantification

Co-staining for laminin-α2 and MyHC-emb was performed on transverse TA sections as previously described. Images were obtained with the Axioplan 2 Microscope System (Carl Zeiss, Germany). Laminin-α2 was used as a mask to identify total fibre number per image. The proportion of MyHC-emb–positive fibres was calculated as the number of MYH3-positive fibres divided by the total number of fibres. Quantification was performed on multiple images covering the entire muscle, and all counting was performed in a blinded manner.

### Protein analyses

Muscle samples were homogenized (Precellys 24, Bertin Technologies) for 2 × 30 s at 5500 rpm on ice in RIPA buffer (R0278, Sigma-Aldrich) supplemented with protease inhibitors (1:100, P8340, Sigma-Aldrich). Protein concentration was determined using a BCA assay (23227, ThermoFisher Scientific). Equal amounts (30 µg) of total protein were heat-denatured (5 min, 100 °C), separated on NuPAGE 3–8% Tris-Acetate Midi Gels (Novex, Life Technologies), and transferred to PVDF membranes (Millipore). Membranes were briefly stained with Ponceau S (P7170, Sigma) to visualize total protein, blocked for 1 h in Odyssey Blocking Buffer (926-41090, LI-COR), and incubated for 2 h at room temperature with the following primary antibodies: anti-utrophin (1:50, MANCHO3(84A), gift from G.E. Morris), anti-MYH3 (1:100, sc-53091, Santa Cruz Biotechnology), anti-dystrophin (1:200, ab15277, Abcam), anti-α-actinin (1:400, sc-7453, Santa Cruz Biotechnology), anti-GAPDH (1:20 000, MA5-15738, Invitrogen), anti-vinculin (1:10 000, ab129002, Abcam). Infrared fluorescence was detected using the Odyssey Imaging System (LI-COR Biosciences, USA). Dystrophin, utrophin, and MYH3 signal intensities were quantified using Image Studio Lite v5.0 (LI-COR Biosciences).

### RNA analyses

Total RNA was extracted from TA muscles using TRIzol™ reagent (ThermoFisher) following the manufacturer’s protocol. cDNA was synthesized from 500 ng RNA using the QuantiTect Reverse Transcription Kit (205313, Qiagen). Quantitative PCR was performed using the 7500 Fast Real-Time PCR System (Applied Biosystems) and Fast SYBR™ Green Master Mix (4385612, ThermoFisher). Reference gene stability was assessed using the geNORM kit (Primer Design, UK; Cat# ge-SY-12), and results were analysed by the ΔΔCT method. Primer sequences were as follows: *utrn-A* (forward primer 5’ ACGAATTCAGTGACATCATTAAGTCC-3’, reverse primer 5’ATCCATTTGGTAAAGGTTTTCTTCTG-3’), *myh3* (forward primer 5’CTTCACCTCTAGCCGGATGGT-3’, reverse primer 5’AATTGTCAGGAGCCACGAAAAT-3’). mRNA expression levels were normalised to *gapdh* (forward primer 5’GTATGACTCCACTCACGGCAAA-3’, reverse primer 5’GGTCTCGCTCCTGGAAGATG -3’) as reference gene (stability value < 1.5). No-RT and no-template controls were included in each 40-cycle PCR run (Cq values: NTC = undetermined; non-RT = undetermined; ALBh (human albumin) > 35).

### Statistics

Data were analysed using GraphPad Prism 8.4.2 (GraphPad Software, La Jolla, CA). Comparisons among wt, mdx, and treatment groups were performed by one-way ANOVA followed by Tukey’s post-hoc test. Two-group comparisons were performed using unpaired Student’s t-tests (two-tailed), assuming equal or unequal variances as determined by F-test. Comparisons involving contralateral saline- and CTX-treated TA muscles were analysed as paired Student’s t-tests (two-tailed). No outliers were excluded. Data are presented as mean ± SEM, where n indicates the number of independent biological replicates. Statistical significance was defined as p < 0.05 (*), p < 0.01 (**), and p < 0.001 (***).

## Acknowledgements

We acknowledge G.E. Morris (Oswestry, UK) for the MANCHO3 antibody.

---

**S1 Fig.** Temporal expression of loading controls following cardiotoxin injection. (A) Representative immunoblots showing vinculin (VINC), α-actinin, GAPDH, and total protein (Ponceau staining) in individual control and CTX-injected wt TA muscles across the time course of CTX-induced muscle regeneration. (B) Representative immunoblots showing vinculin (VINC), α-actinin, GAPDH, and total protein (Ponceau staining) in individual control and CTX-injected mdx TA muscles across the same time course.

**S2 Fig.** Stability analysis of housekeeping genes. Twelve candidate housekeeping genes were evaluated using all wt and mdx samples across the time course of CTX-induced muscle regeneration to identify the most stable reference gene(s). Gene expression stability was assessed using geNORM.

**S1 Data.** Individual numerical data underlying the graphs and statistical analyses presented in this study.

